# Developing a unique medical-grade honey which maximizes glucose oxidase activity

**DOI:** 10.1101/411819

**Authors:** Zhiyuan Li, Rohan Parikh

**Affiliations:** Loudoun Academy of Science, 21326 Augusta Drive, Sterling VA, 20164 |

**Keywords:** medical-grade honey, glucose oxidase, hydrogen peroxide, antimicrobial agents, anticancer agents

## Abstract

In the fight against cancer and infection, honey is a compelling solution by virtue of its unique chemical composition; however, current medical-grade honeys are expensive and limited in their scope. In this study, a novel medical-grade honey was developed by maximizing the activity of glucose oxidase (GOX), an enzyme in honey that synthesizes hydrogen peroxide; maximization was done by neutralizing the effects of catalase and methylglyoxal (MGO), compounds in honey that interfere with H_2_O_2_ accumulation. Expressed GOX activity was quantified via H_2_O_2_ accumulation in honey after catalase, MGO, or both were inhibited. Results indicate the honeys tested have significant quantities of H_2_O_2_ inhibitors, greatly affecting expressed GOX activity; neutralization resulted in at least a 100% increase in H_2_O_2_ accumulation in all honeys. Blueberry honey with catalase inhibition by EGCG saw a 938% increase in H_2_O_2_ accumulation, reaching nearly three times the H_2_O_2_ accumulation in current medical grade honey for a tenth of the cost. This research presents a novel and readily reproducible method for maximizing H_2_O_2_ accumulation in any honey through neutralization of GOX inhibitors. GOX activity enhancement, combined with honey’s diverse antioxidants, enables the emergence of global low-cost medical-grade honeys with immense potential to revolutionize cancer and infection treatment.

This paper is based on work presented at the Regional Science and Engineering Fair on March 15, 2018 at Riverside High School (Loudoun County, Virginia), the Virginia State Science and Engineering Fair on April 14, 2018 at the Virginia Tech Carilion School of Medicine (Roanoke, Virginia).

**Author Summary:** With cancer and post-operative infections on the rise, a natural, low-cost alternative treatment is necessary. Honey is an ideal candidate due to its diverse antioxidants and glucose oxidase, an enzyme that produces hydrogen peroxide. The purpose of this research was to maximize glucose oxidase activity by neutralizing compounds that interfere with H_2_O_2_ accumulation. This research yielded medical-grade honeys that are economical and more effective than current ones, presenting a novel method that is reproducible on a large scale, scalable to different needs, and applicable to any honey to create low-cost medical-grade honeys that can revolutionize cancer and infection treatment globally.

## Introduction

Cancer and infections are two of the most pressing issues facing the world; infections and related complications are incredibly common in post-surgical recovery, and cancer is extremely widespread, in both developed and developing nations [1]. Current treatments for infections include antibiotics and medical dressings; however, microbial resistance is becoming increasingly common, as diseases evolve to resist antibiotic treatments [2]. Current treatments for cancer include chemotherapy and radiation therapy; however, treatment can last anywhere from 3 to 6 months, and the immune system remains immunocompromised for 21 to 28 days after treatment [3].

The condition of post-operative care in developing nations, such as in Africa, make patients twice as likely to die after surgery due to complications and infections. In addition, the lack of certainty and degradation of quality of life that current cancer treatments offer make them unfeasible for developing nations, as they generally do not have the money or ability to pay for the treatments and care for their patients [4]. Therefore, a low-cost, readily accessible, and safer alternative is needed to combat cancer and infections, both in developing nations and locations such as the United States; honey presents itself as an ideal candidate for this alternative.

Honey is a compelling anticancer agent by virtue of its unique chemical composition. It provides a protective barrier, maintains a moist wound condition, and promotes healing by stimulating growth and migration of new cells and essential proteins to wound sites. Its low pH is acidic enough to inhibit the growth of most bacteria, while high sugar content honey can physically dehydrate microbes [5][6][7]. Antioxidants in honey inhibit tumor growth, reduce inflammation, minimize cytotoxic effects, and can induce programmed cell death in cancer cells [8]. Furthermore, studies have shown that honey has the ability to reduce infections in throat-cancer patients; honey was applied to throat cancer sores and stopped the progression of infection completely, while control groups without the applied honey exhibited progression of infection to Grade 3 and 4 [9]. Thus, even if medical grade honey cannot be used as a primary cancer treatment, it can be used in conjunction with current treatments to alleviate side effects and minimize cytotoxic fallout.

There are two types of current medical grade honey: peroxidase and non-peroxidase. The first and most prevalent form of medical-grade honey is manuka honey. This honey is unique in that it has an overwhelming presence of methylglyoxal (MGO), a non-enzymatically formed compound derived from flower nectar that exists in varying quantities within honeys [10]. MGO has been shown to have antibacterial properties against both gram-positive and gram-negative bacteria, as well as multi-resistant strains when in high concentrations; this antibacterial activity is resistant to heat and light degradation. However, a major problem with manuka honey is that it is a potential risk factor in healing diabetic ulcers and can increase the risk of infection in diabetic patients after amputation when applied topically in conjunction with medical dressings [10]. This is especially an issue in patients with peripheral artery disease (PAD), as this condition is particularly prevalent in type 2 diabetic patients and can result in amputation [11]. The mechanism by which MGO is a risk factor for diabetics is MGO glycation, the formation of adducts to blood sugars, putting diabetic patients at risk [10]. Furthermore, MGO is a reactive metabolite that can exert toxic effects; while MGO levels in manuka honey are typically safe, batches of manuka honey are increasingly selected for maximized levels of MGO, raising toxicity concerns which need to be investigated [12].

The other type of medical-grade honey is peroxidase-based; the honey utilizes the accumulation of hydrogen peroxide (H_2_O_2_) as its primary antibacterial component. Most honeys are peroxidase; however, the H_2_O_2_ accumulation in most natural honeys is too low to be particularly effective, and therefore these honeys do not lend well to medical applications [5]. The primary peroxidase medical-grade honey is Revamil (RS) honey, a commercially available medical-grade honey that has a very low MGO level and high H_2_O_2_ accumulation, which is synthesized by an enzyme, glucose oxidase (GOX). GOX is synthesized in the hypopharyngeal gland of bees to sterilize colony food, and facilitates the formation of H_2_O_2_, which contributes to honey’s antibacterial activity, and gluconic acid, which lowers honey’s pH, from glucose, water, and oxygen [13]. Maximization of its expressed activity in honey is essential to creating an effective peroxidase medical-grade honey. RS honey does this by selectively breeding flowers in a greenhouse in the Netherlands; the resulting honey accumulates as much as 4.4 mM of H_2_O_2_ [12][14].

While these honeys are effective as medical-grade honeys, one major problem of these honeys is their expense; 15 grams of the RS honey costs $25, and similar costs are seen for manuka honey, rendering them unrealistic options for large-scale consumption and distribution, especially in developing nations. This expense is attributed to the aforementioned selective breeding, which requires many resources. In the face of these unrealistic and limited medical-grade honeys, an alternative low-cost, equally effective medical-grade honey is vital.

There are two compounds also present in honey that interfere with GOX activity in most standard raw honeys: catalase and MGO [15][7]. Catalase is a ubiquitous enzyme, also found in human cells, that catalyzes the decomposition of H_2_O_2_ into water and oxygen, thus interfering with H_2_O_2_ accumulation. Previous literature has identified its presence in honey, and has shown that it apparently regulates the activity of GOX by controlling the H_2_O_2_ equilibrium [15]. MGO, previously mentioned due to its antibacterial effects in manuka honey, also inhibits GOX via the formation of high molecular weight adducts, thus changing enzymatic structure and interfering with activity [7].

The purpose of this research is to mimic RS honey in its mechanism of antibacterial activity, through H_2_O_2_ accumulation; however, instead of selective breeding, this research uses the method of neutralizing the inhibitors of GOX in honey (catalase and MGO), and thus maximize the expressed GOX activity. MGO is neutralized via the glyoxalase system [12]. Catalase can be neutralized by two separate compounds,-mercaptoethanol and epigallocatechin-gallate (EGCG). ß-mercaptoethanol is a chemical commonly used as a reducing agent, and has been shown to inhibit catalase by reducing disulfide bonds in the tertiary structure of the enzyme [16]. Meanwhile, EGCG is a catechin found in the green tea leaf plant, and has various anti-cancer and antioxidant properties. It has been shown to partially inhibit catalase solutions as a noncompetitive inhibitor at certain concentrations, but its ability to fully inhibit catalase has not been observed [17]. The effect of these inhibitors on GOX is unknown. Prior research on the structure of GOX has found that the disulfide bonds in GOX are hidden inside the protein, and are therefore not available for reaction unless the enzyme is unraveled (denatured) through heating. Because of the placement of the disulfide bonds, it was hypothesized that ß-mercaptoethanol would not be able to reduce the disulfide bonds in GOX and denature it. Regarding EGCG, however, prior research cannot conclusively state its effects on GOX.

It was hypothesized that the neutralization of catalase and MGO would increase expressed GOX activity regardless of floral source because the two compounds interfere with GOX by decomposing H_2_O_2_ and forming adducts to change the enzymatic structure. Additionally, it was hypothesized that a greater expressed GOX activity in a honey would correlate to higher GOX content due to the direct, positive relationship between enzyme concentration and rate of reaction in a system with unlimited substrate.

The presence of the inhibitors, catalase and MGO, was varied so that each were neutralized separately and together in a honey sample. In addition, the floral source was varied. The dependent variable was expressed GOX activity, quantified by H_2_O_2_ accumulation. The control group for each honey consisted of raw, sterile, diluted honey; raw honey was necessary in order to preserve the enzymes that are normally denatured in pasteurized (heated) honey. Sterile honey was required due to the presence of microorganisms in honey such as bacteria and yeast, which are activated and begin proliferation once water is added; water needed to be added to dilute the honey in order to activate the GOX enzyme and reduce its viscosity for sterilization.

## Materials & Methods

Four honeys were used in this investigation; of the four honeys tested, two were monofloral honeys, or honeys that are derived primarily from one species of flower. The other two were polyfloral honeys, honeys derived from multiple flowers with no primary flower type. The monofloral varieties were blueberry and clover, while the polyfloral honeys were wildflower honeys from Ohio and Pennsylvania - all honeys were from the United States.

Group A is used to denote the control group, in which catalase and MGO were both active and no additional chemicals will be added. Group B is used to denote catalase inhibition, in which catalase is inhibited but MGO is active, in order to test the effect of catalase on GOX. Group C is used to denote MGO inhibition, in which MGO is inhibited but catalase is active, in order to test the effect of MGO on GOX. Finally, Group D is used to denote combined neutralization, in which neither catalase nor MGO are active, to maximize GOX to its full potential.

10 grams of honey were diluted with 10 mL of diH_2_O. The honey solution was filtered through a 0.22 μm micron-filter (Millipore; Product #: SVGP01015) in a sterile Büchner funnel. The funnel and flask were wrapped in aluminum foil to protect the honey solution from light. The aforementioned steps were repeated as necessary to sterilize enough honey for Groups A - D. Any further reference to “Group A” or “honey solution” refers to sterilized, raw, diluted honey.

Pre-testing procedures were done for catalase inhibition to determine the necessary concentrations and incubation time (pre-testing was not done for MGO inhibition because the procedure in honey is established). Catalase was inhibited with two different compounds: ß-mercaptoethanol (ß-met) (Sigma Aldrich; Product #: M6250) and epigallocatechin gallate (EGCG) (Sigma Aldrich; Product #: E4268). To inhibit catalase, ß-met was added to a final concentration of 5 mM in a fume hood (3.5 L ß–met/10 mL of honey solution); this concentration was based on established literature that 5 mM of ß–met inhibits 50% catalase activity after 1 hour of incubation [16]. The procedure for EGCG was done by adding EGCG in three different concentrations (100 μM, 200 μM, 400 μM). Concentrations were initially based on established research that used 50 μM EGCG to inhibit catalase activity after 1 hour of catalase; however, this concentration did not work for the honey samples, so the concentrations and incubation time were increased [17].

After 48 hours of incubation, a visual catalase assay was conducted on the ß-met and EGCG-treated honey solutions to determine whether the enzyme is completely inhibited; equal volumes of 30% H_2_O_2_ were added to each treated honey solution and the respective untreated honey, and the formation or lack of formation of bubbles was observed.

Complete inhibition was seen with the 5 mM of ß-met with 48 hours of incubation time and 400 μM of EGCG with 48 hours of incubation time. These values were used to prepare Groups A - D; EGCG inhibition was not done from the very beginning of the activity tests due to time constraint, and because it was not established as a catalase inhibitor in previous literature, while ß-met is more reliable as an established, though hazardous, inhibitor.

Group A was the control group, Group B had catalase inhibited, the extension of Group B had catalase inhibited with EGCG, Group C had MGO inhibited via the glyoxalase system using reduced glutathione (Sigma Aldrich; Product #: G4251) and the glyoxalase-I enzyme (Sigma Aldrich; Product #: G4252). Glyoxalase-I was added in a 1:2 UN of enzyme to mL of honey ratio and incubated for 30 minutes. Group D was prepared by combining the glyoxalase system and the ß-met.

A fluorescence assay to quantify expressed GOX activity by H_2_O_2_ accumulation was run on a Gen 5 Synergy HT microplate reader (Bio-Tek; Part #: 70910000) as per assay specifications of the Amplex Red Glucose/Glucose Oxidase Assay Kit (Thermofisher Scientific; Product #: A22189) for all Groups A - D (each group had 4 honeys, the two monofloral honeys and two polyfloral honeys). One change that was made to the assay directions was that the solution that was put in the well plate was 50 μL, but broken down as 25 μL honey solution and 25 μL 1X reaction buffer.

After activity testing was completed, qualitative observations of protein content in the honey were taken using an SDS-PAGE procedure. However, the procedure could not be done with pure honey; thus, the proteins were extracted from two honeys. The two honeys used were the blueberry and clover honey; these two were chosen because blueberry had the greatest expressed GOX activity, while clover had the least. Therefore, differences in protein content to support the hypothesis that greater activity correlated to greater content would be the most easily observable. 20 grams of raw honey was measured out and diluted with 20 mL of dH_2_O. One 30 cm strip of dialysis tubing with molecular weight cutoff of 14 kDa (Sargent Welch; Product #: 470163) was used to dialyze the honey solution against 2 L of dH_2_O and stirred continuously on a stir plate (Phenix; Product #: H4000-HS) for 48 hours.

The contents inside the dialysis tubing were concentrated via centrifugation (5000 rpm for 10 minutes at a time). The solution was quantified using a Pearl nanophotometer (Implen; Product #: P300); the SDS-PAGE samples were run at a concentration of ~2000 μg/mL. Two SDS-PAGE gels were run in order to observe the GOX protein; established research showed GOX in honey was approximately 120 kDa, and was a dimer; thus, when linearized via SDS-PAGE, protein bands would be seen around 50-60 kDa. The SDS-PAGE procedure was run using two gel electrophoresis chambers (Edvotek Inc.; Product #: S90257ND), in which 7.5% Mini-Protean TGX Precast 12-well, 20 μL polyacrylamide gels (Bio-Rad; #4561025dc) were used. Tris-glycine-SDS 1X running buffer (10X concentrate from Bio-Rad; Product #: 161-0732) was used to run the gel. The gels were run at 100 V and then transferred.

The gels were transferred into AcquaStain Protein Gel Stain (Bulldog Bio; Product #AS001000) and were stained for 48 hours on a rocker. LoggerPro gel analysis software (Vernier; Product #: LP) was used to quantify the size of the proteins observed on the stained gel.

## Discussion and Conclusions

Kruskal-Wallis’, ANOVAs, Mann-Whitneys, t-tests, and a 2-way ANOVA with replication tests were performed. The difference between each of the honeys, control and experimental groups, and pollen variety groups as a whole was determined. P-values as low as 7 x 10-28 (alpha = 0.05) in the 2-way ANOVA test imply that there was significant difference between honeys, combinations of neutralization, and their combined effects. In particular, there was a statistically significant increase in the expressed GOX activity in one or more of the experimental groups when compared to the control for every single honey tested. This provides support for the assertion that the catalase and MGO were interfering with the expressed GOX activity, regardless of content of catalase, MGO, or the baseline GOX activity.

Literature on Revamil (RS) honey has shown that maximum H_2_O_2_ accumulation can reach 4.4 mM final concentration [12][14]. Prior to the addition of any inhibitors, the honeys used in the trial reached, on average, 20% of that maximum RS honey accumulation, with a lowest in clover honey at 8.11% and a highest in WF 1 at 32.65% (Table 1). After catalase was inhibited, all honeys saw at least a 100% increase in expressed GOX activity (Table 1). The most marked increases in activity was in blueberry honey after catalase inhibition and WF2 after MGO neutralization. In WF2, the findings show an extremely significant increase in expressed GOX activity by 366% after MGO was inhibited compared to the control (P < 0.001) (Table 1). Regarding blueberry honey, there was an extremely significant increase in expressed GOX activity by 574% after catalase was inhibited (P < 0.001); blueberry honey reached the greatest H_2_O_2_ accumulation out of all four honeys tested (Table 1).

**Table 1.** Hydrogen peroxide accumulation (mean ± standard deviation and range) in the four tested honeys. Group A was control honey (untreated), Group B had catalase inhibited with beta-mercaptoethanol, Group C had MGO inhibited with the glyoxalase system (reduced glutathione and glyoxalase I), while Group D had combined inhibition of catalase and methylglyoxal. Group B with EGCG had catalase inhibited with epigallocatechin gallate instead of beta-mercaptoethanol and was only tested for blueberry honey.

| Group | Blueberry H<sub>2</sub>O<sub>2</sub> accumulation (mM) |  | Clover H<sub>2</sub>O<sub>2</sub> accumulation (mM) |  | Pennsylvania Wildflower H<sub>2</sub>O<sub>2</sub> accumulation (mM) |  | Ohio Wildflower H<sub>2</sub>O<sub>2</sub> accumulation (mM) |  |
| --- | --- | --- | --- | --- | --- | --- | --- | --- |
|  | Mean | Range | Mean | Range | Mean | Range | Mean | Range |
| **A** | 0.85 ± 0.21 | 0.51 - 1.12 | 0.55 ± 0.10 | 0.41 - 0.72 | 1.34 ± 0.17 | 1.04 - 1.63 | 0.99 ± 0.19 | 0.59 - 1.20 |
| **B** | 5.52 ± 0.45 | 5.14 - 6.31 | 1.13 ± 0.25 | 0.85 - 1.48 | 2.69 ± 0.15 | 2.52 - 2.96 | 2.88 ± 0.14 | 2.68 - 3.15 |
| **B with EGCG** | 8.86 ± 1.66 | 6.70 - 11.14 | ---- | | | | | |
| **C** | 3.56 ± 0.57 | 3.00 - 4.54 | 1.39 ± 0.09 | 1.29 - 1.56 | 1.88 ± 0.10 | 1.72 - 2.02 | 4.63 ± 0.43 | 4.18 - 5.28 |
| **D** | 2.95 ± 0.16 | 2.62 - 3.24 | 1.39 ± 0.11 | 1.25 - 1.56 | 1.96 ± 0.38 | 1.70 - 2.95 | 2.73 ± 0.27 | 2.41 - 3.16 |

Further, using a less harmful catalase inhibitor during EGCG post-testing yielded an even greater increase in activity than the ß-mercaptoethanol, which demonstrates that EGCG can enhance GOX activity. Moreover, it implies that the ß-mercaptoethanol, which is a known reducer of disulfide bonds common to enzymes and proteins, affected the activity of GOX in the originally designed catalase inhibition trials, despite the presumed location of the disulfide bonds as hidden within the enzyme. Statistically, adding EGCG rather than beta-mercaptoethanol to blueberry honey showed an extremely significant increase in expressed GOX activity by 60% (p < 0.0001); the resulting H_2_O_2_ accumulation showed a 938% increase from the control blueberry honey (Table 1, Figure 2). When compared to RS honey, its maximum H_2_O_2_ accumulation was 11.14 mM with EGCG, nearly three times the maximum accumulation in RS honey.

**Figure 1.**
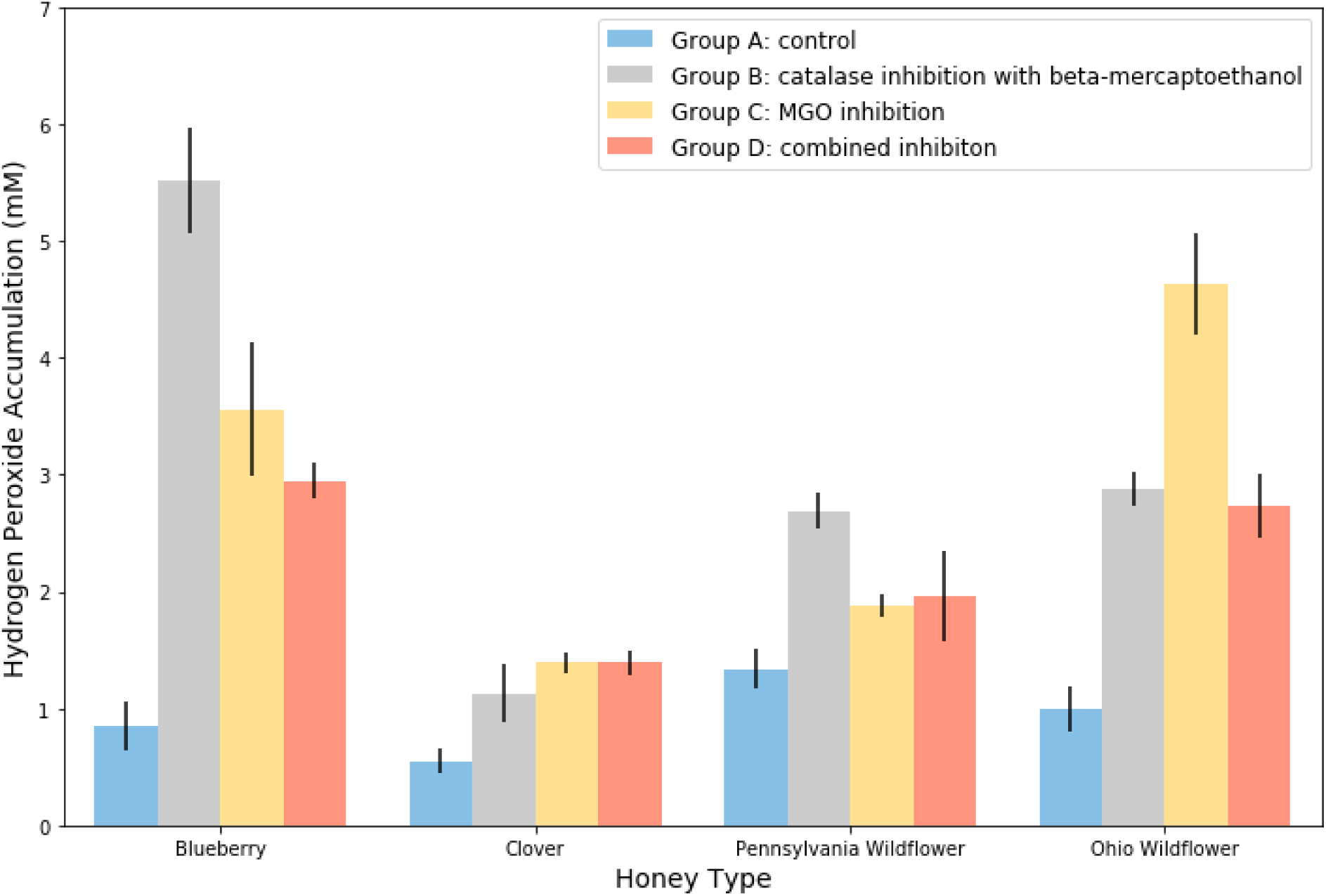
H_2_O_2_ accumulation in mM, indicative of GOX activity, as a function of experimental group and honey. Error bars represent standard deviation.

**Figure 2.**
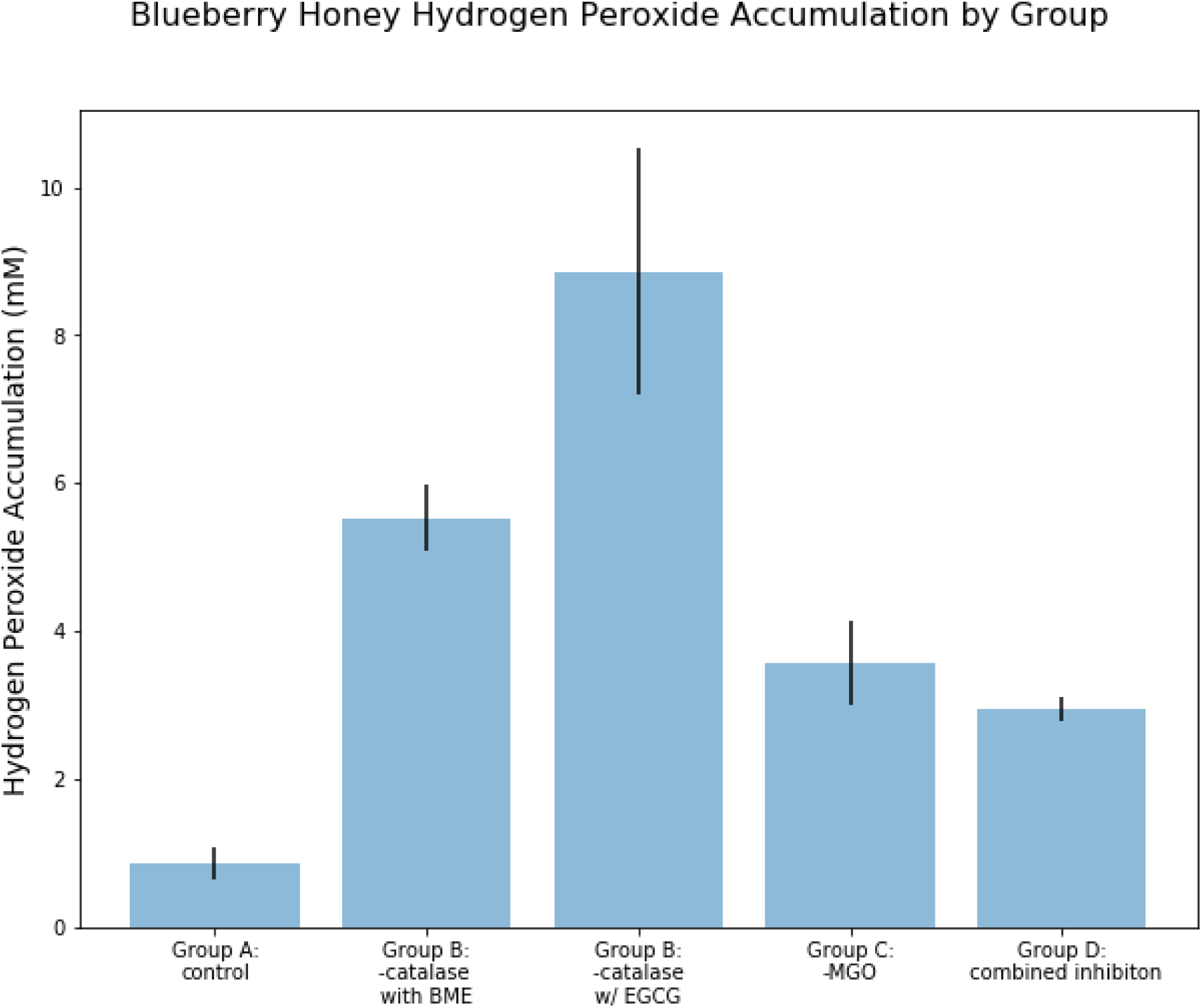
H_2_O_2_ accumulation in mM, indicative of GOX activity, as a function of experimental group for blueberry honey with the Group B extension included. Error bars represent standard deviation.

**Figure 3.**
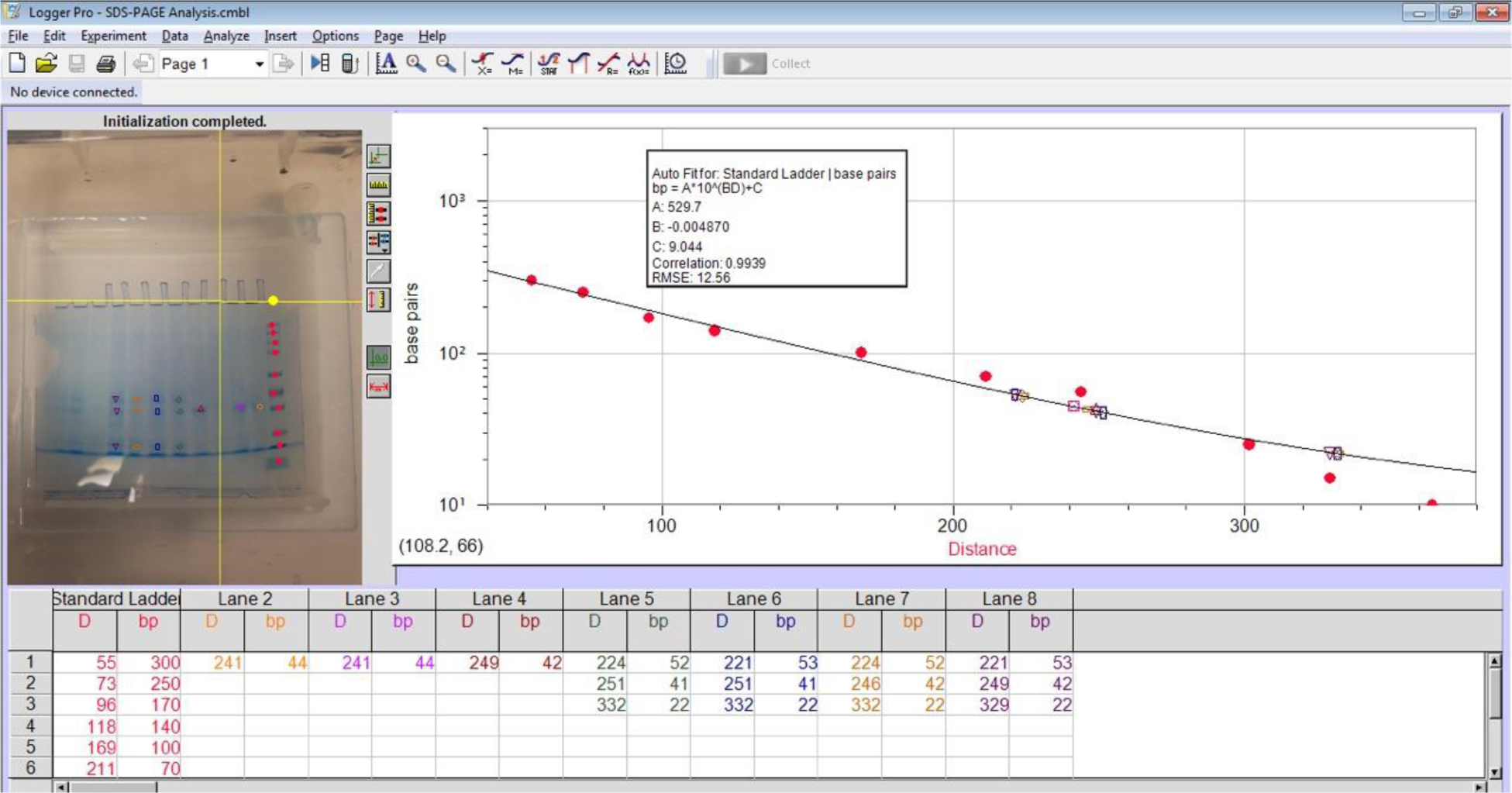
Protein gel analysis software and standard curve generated from protein standard. The picture of the gel was analyzed, with the distinct bands marked in the software.

Interestingly, Group D saw a significant drop in expressed GOX activity when both catalase and MGO were neutralized, rather than showing the greatest increase in activity due to a synergistic effect of both inhibitors (Table 1). This is likely attributed to the mechanism of inhibition by the ß-met that interfered with the MGO inhibitor, the glyoxalase system. MGO is inhibited by an enzyme in the presence of reduced glutathione by conversion to another compound, while ß-met reduces disulfide bonds, which are extremely common in enzymes. It is feasible that the ß–met reduced disulfide bonds in the glyoxalase-I enzyme as well, rendering it ineffective, and preventing the ß–met from neutralizing catalase; future work would include using an alternate inhibitor of catalase in conjunction with the glyoxalase system to further maximize GOX activity in Group D.

A major problem with current medical-grade honeys is their extremely high costs due to expensive selective breeding. Thus, cost-benefit analysis was essential in order to compare both the effectiveness and expense of an alternative honey to RS honey. The cost of the blueberry honey, which had the highest expressed GOX activity, was combined with the cost of the EGCG tea catechin and the cost of the glyoxalase system, under the assumption that future work would maximize GOX activity further. The costs also incorporated shipping and handling and did not account for lower wholesale costs. Thus, the exceedingly liberal estimate of the maximum cost of the alternative medical-grade honey was $2.78 for 15 grams, in comparison to $25 for 15 grams of RS honey. The alternative presented was therefore 3 times as effective as RS honey, with 11.14 mM of H_2_O_2_ accumulation compared to 4.4 mM, at merely a tenth of the cost (See Appendix B for full cost-benefit analysis).

Some additional conclusions were made by virtue of the methodology conducted in this research. EGCG has been identified as a partial catalase inhibitor, but full inhibition has not been observed [17]. In addition, established research has not investigated the effect of EGCG on GOX. This research presents EGCG as a complete, non-competitive inhibitor of catalase that also does not inhibit GOX and may even enhance its activity. This is corroborated by the increased levels of expressed GOX activity in EGCG-treated blueberry honey. Another explanation is that ß-met did actually inhibit GOX to some degree, but only mildly, seen by the relatively high levels of H_2_O_2_ in ß-met-treated honey; thus, EGCG-treated honey, which had higher levels of GOX activity, may enhance GOX but does not inhibit the enzyme at all.

The enzymatic structure of GOX has been observed in fungi and yeast, such as *Aspergillus niger*, the structure of GOX in honey has not been identified. However, the statistically significant difference of an over 100% increase with catalase inhibition via ß–met, which reduced catalase’s disulfide bonds, gives clues into the structure of GOX in honey; most or all of the disulfide bonds are protected from inhibition, likely due to their location in the center of the enzyme. In addition, catalase has been identified in honey as an inhibitor of GOX, but recent research on the antibacterial activity of honey have failed to consider its significance in regulating H_2_O_2_ accumulation in honey. By neutralizing catalase on its own in Group B, the varying amounts of catalase in multiple honeys was observed, as well as its significance in controlling expressed GOX activity. It is seen by this methodology that, despite certain similar levels of baseline GOX activity, there are varying quantities of both MGO and catalase in honey that correspond to different maximum levels of expressed GOX activity.

Protein analysis was conducted in order to test the latter part of the hypothesis, that greater GOX activity would correspond to greater GOX content. The gel electrophoresis was conducted for the blueberry and clover honeys and LoggerPro software was used to identify the visible bands on the polyacrylamide gel; bands of ~50-60 kDa were most likely GOX because GOX in honey is ~120 kDa and a linearized dimer [18]. The only distinct bands visible on the gel for the lanes in which clover samples were run were at ~42 - 44 kDa, which was hypothesized to be catalase (160 kDa with 4 identical subunits that were linearized) (Table 2). However, there were distinct bands visible for the lanes in which blueberry samples were run at 52-53 kDa, which were hypothesized to be GOX (Table 3). This hypothesis stems from a qualitative observation and could not be confirmed without a Western blot test with anti-glucose oxidase antibodies; however, the sizes did correlate fairly well. This data partially supports the hypothesis that greater activity corresponds to greater content, as bands of protein which was potentially GOX were observed in the blueberry honey, which had the highest activity, but were not observed in the clover honey, which had the least activity.

**Table 2.**
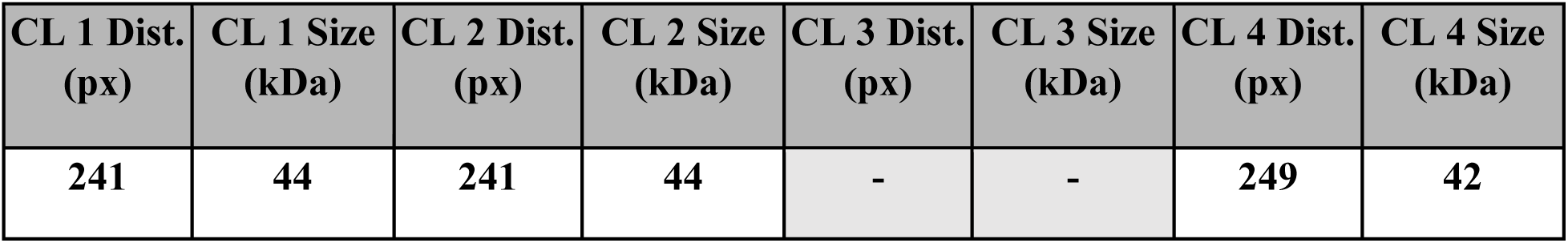
Protein bands found in the clover (CL) honey samples. Using the Logger Pro gel analysis software, a picture of the gel was analyzed; the distance from the bottom of the well (in pixels) was correlated to protein size in distinct bands using a standard curve generated from the concurrently run protein standard. Distinct bands of interest could only be found in the 42-44 kDa range in samples 1, 2, and 4; in sample 3, no bands were found that could possibly be glucose oxidase.

**Table 3.** Protein bands found in the blueberry (BB) honey samples. Using the Logger Pro gel analysis software, a picture of the gel was analyzed; the distance from the bottom of the well (in pixels) was correlated to protein size in distinct bands using a standard curve generated from the concurrently run protein standard.

| BB 1 Dist.<br>(px) | BB 1 Size<br>(kDa) | BB 2 Dist.<br>(px) | BB 2 Size<br>(kDa) | BB 3 Dist.<br>(px) | BB 3 Size<br>(kDa) | BB 4 Dist.<br>(px) | BB 4 Size<br>(kDa) |
| --- | --- | --- | --- | --- | --- | --- | --- |
| 224 | 52 | 221 | 53 | 224 | 52 | 221 | 53 |
| 251 | 41 | 251 | 41 | 246 | 42 | 249 | 42 |
| 332 | 22 | 332 | 22 | 332 | 22 | 329 | 22 |

While the methodology maximizes GOX activity to a large extent, there are several factors that can be adjusted in order to further optimize antibacterial activity. While we used the same dilution for all of our honeys, different honeys have different optimal dilutions for maximum H_2_O_2_ accumulation. Thus, an optimal dilution could be found by varying dilution of honey to 1X reaction buffer. The dilution used in this investigation was constant across honeys strictly for the purpose of comparison, but this can easily be changed to allow for further maximization. In addition, in Group D, the combined neutralization, it was hypothesized that the H_2_O_2_ would be maximized the most; however, there was a statistically significant drop in activity when both catalase and MGO were neutralized, likely due to interference of the inhibiting compounds. Thus, to maximize expressed GOX activity even further, this group could be tested again using EGCG and the glyoxalase system rather than ß-met. Lastly, the content of GOX can be quantified using a western blot protein quantification method in order to compare enzyme content with respect to the variation in expressed GOX activity data seen in this research.

Current medical-grade honey can be significantly advanced by the development of a low-cost alternative; this research presents such an alternative, with the promising candidate of EGCG-treated blueberry honey, which exhibited greater effectiveness than current medical-grade honey for a mere fraction of the cost. However, the development of just one alternative low-cost honey, while significant, does not address the overarching problem concerning a limited resource pool and widespread circulation for medical-grade honeys. Developing a method that can make any honey medical-grade broadens the feasible resource pool for application in the medical field, thus relieving the need for large-scale distribution. Thus, this research establishes a completely novel and readily reproducible method for maximizing H_2_O_2_ accumulation in any honey through the neutralization of GOX inhibitors. GOX activity enhancement, combined with honey’s diverse antioxidants, enables the emergence of low-cost medical-grade honeys possessing immense potential to revolutionize cancer and infection treatment in developing nations.

## Acknowledgements

The authors received funding from the Loudoun Academy of Science; no competing interests exist. The authors sincerely thank Ms. Jackie Curley of the Loudoun Academy of Science for providing the guidance and materials needed to conduct this research; your passion and dedication as a mentor inspired us to develop our own paths to discovery.

